# Triose phosphate utilization and beyond: from photosynthesis to end-product synthesis

**DOI:** 10.1101/434928

**Authors:** Alan M. McClain, Thomas D. Sharkey

## Abstract

During photosynthesis plants fix CO_2_ from the atmosphere onto ribulose-bisphosphate producing 3-phosphoglycerate, which is reduced to triose phosphates (TPs). The TPs are then converted into the end products of photosynthesis. When a plant is photosynthesizing very quickly it may not be possible to commit photosynthate to end product as fast as it is produced, causing a decrease in available phosphate and limiting the rate of photosynthesis to the rate of triose phosphate utilization (TPU). The occurrence of an observable TPU limitation is highly variable based on species and especially growth conditions, with TPU capacity seemingly regulated to be in just slight excess of the likely photosynthetic rate. The physiological effects of TPU limitation are discussed with an emphasis on interactions between the Calvin-Benson cycle and the light reactions. Methods for detecting TPU-limited data from gas exchange data are detailed, and the impact on modeling of some physiological effects are shown. Special consideration is given to common misconceptions about TPU.

**Highlight:** Photosynthetic triose phosphate utilization limitation is discussed, highlighting misleading points in physiology and focusing on regulation.

## Introduction

Triose phosphate utilization (TPU) is one of the three canonical biochemical limitations of photosynthesis in gas exchange analysis of C_3_ plants. It reflects a steady-state condition in which assimilation of carbon is limited by the ability to regenerate phosphate through production of end-products of photosynthesis. Phosphate is required by ATP synthase to produce ATP, of which three are needed to fix a single carbon. Although all three ATP are used for phosphorylation of carbon chains, two are immediately released when the 3-phosphoglyceric acid (PGA) kinase reaction is followed by glyceraldehyde 3-phosphate (GAP) dehydrogenase. One phosphate per three carbons is carried through to the triose phosphates (TPs) glyceraldehyde 3-phosphate and dihydroxyacetone phosphate (DHAP). If the plant is in favorable enough conditions and is photosynthesizing very fast, it is possible to fix three carbons faster than they can be processed to end products. In this scenario, the phosphate concentration decreases, leading to reduced conductivity of protons through thylakoid ATP synthase that ultimately slows photosynthesis (Kanazawa and Kramer, 2002; Takizawa *et al.*, 2008; Kiirats *et al.*, 2009). This is a form of feedback limitation and is potentially quite dangerous to the plant. Feedback conditions are known to cause photodamage due to the inability to move energy downstream (Pammenter *et al.*, 1993; Takizawa *et al.*, 2008; Kiirats *et al.*, 2009). To avoid photodamage, instead of maintaining phosphate-restricted feedback, a series of regulatory steps are engaged to slow photosynthesis down. While the rate is determined by phosphate balance, steady-state rate is set by regulatory effects that serve to ameliorate feedback conditions. This includes reduction in the photosystem 2 quantum yield (*Φ*_*PS2*_) (Sharkey *et al.*, 1988) and reduced activation state of rubisco (Sharkey *et al.*, 1986a; Viil *et al.*, 2004; Cen and Sage, 2005). In this review we discuss the effect of end-product synthesis on the overall rate and regulation of photosynthesis.

## How are triose phosphates used?

The maximal photosynthetic rate under TPU is primarily, but not exclusively, determined by the rate of conversion of triose phosphates into starch and sucrose. The limitation on assimilation is based on the release of phosphate from Calvin-Benson cycle intermediates, and the most immediate release is from the activity of fructose-1,6-bisphosphatase (FBPase) in the chloroplast or cytosol. Sucrose synthesis occurs in the cytosol, beginning with the translocation of TPs through the triose phosphate/phosphate antiporter (TPT) (Riesmeier *et al.*, 1993). This removes carbon from the Calvin-Benson cycle and returns phosphate from the cytosol to the chloroplast. Each sucrose requires the combination of two hexose molecules, for a total of four triose phosphates. Net phosphate release from organic phosphates in the cytosol occurs at FBPase, UDP-glucose pyrophosphorylase, and sucrose-phosphate phosphatase. Sucrose synthesis has been measured at between 35 and 50% of total carbon assimilation (Sharkey *et al.*, 1985; Escobar-Gutiérrez and Gaudillère, 1997; Abadie *et al.*, 2018). In starch synthesis phosphate release occurs at stromal FBPase and ADP-glucose pyrophosphorylase. The rate of flux through starch varies considerably with the growth conditions of the plant, with *Arabidopsis* growing in an 18 h photoperiod committing only 24% of fixed carbon to starch but when grown in a 6 h photoperiod committing 51% (Sulpice *et al.*, 2014). Other studies show between 30-55% of fixed carbon goes to starch (Sharkey *et al.*, 1985; Escobar-Gutiérrez and Gaudillère, 1997; Abadie *et al.*, 2018). A small amount of phosphate is added to starch in photosynthesizing leaves by glucan-water dikinase and phosphoglucan-water dikinase but the amount is very low, 0.1-0.9% (McPherson and Jane, 1999; Ritte *et al.*, 2002; Kötting *et al.*, 2004), and so is not relevant for discussing gas exchange properties of photosynthesis.

There are a number of other routes by which carbon is exported from the Calvin-Benson cycle (Fig. 1). Any carbon metabolism pathway that begins with a phosphorylated Calvin-Benson cycle intermediate and ends with a non-phosphorylated molecule will contribute to TPU. The shikimate pathway to aromatic amino acid synthesis begins with the export of GAP from the chloroplast to make phosphoenolpyruvate (PEP). PEP is reimported through the phosphoenolpyruvate/phosphate translocator (PPT) and combines with erythrose-4-phosphate (E4P) and ends with chorismate, accounting for 1-2% of photosynthesis (Escobar-Gutiérrez and Gaudillère, 1997; Abadie *et al.*, 2018). Lipids and branched chain amino acids are synthesized from acetyl-CoA from pyruvate and from PEP, accounting for 1-3% of fixed carbon (Bao *et al.*, 2000). It has been shown to be possible to increase oil biosynthesis as a carbon sink, and this would contribute to a higher capacity for TPU (Sanjaya *et al.*, 2011). The methylerithritol phosphate (MEP) pathway begins with GAP and pyruvate to produce isoprenoids, up to 3% of assimilation (Rasulov *et al.*, 2014). Amino acid intermediates in the photorespiratory pathway can be used in the cytosol for protein construction, from an average of 30% to a high of 70% of photorespiration (Busch and Sage, 2017). If the ratio of oxygenation to carboxylation (*φ*) is assumed to be 0.25, this represents 7-15% of photosynthetic carbon.

**Fig 1.**
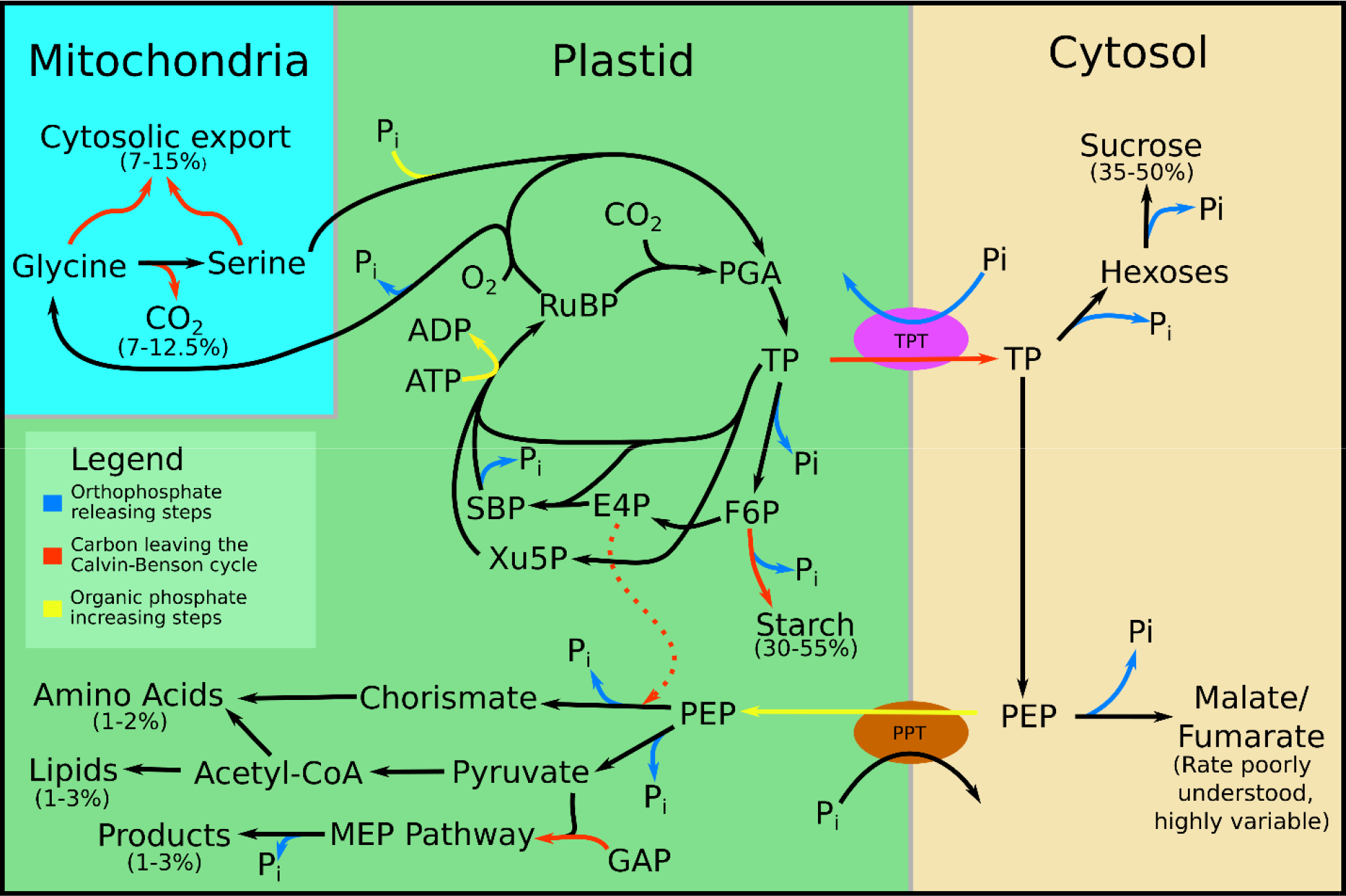
A depiction of the major phosphate and carbon exits from the Calvin-Benson cycle. Abbreviations: E4P, erythrose 4-phosphate; F6P, fructose 6-phosphate; GAP, glyceraldehyde 3-phosphate; PEP, phospho*enol*pyruvate; PGA, 3-phosphoglyceric acid; SBP, sedoheptulose bisphosphate; TP, triose phosphates; Xu5P, xylulose 5-phosphate

Plants are capable of carboxylating PEP then reducing the oxaloacetate to malate for use in anapleurotic reactions or storage in the vacuole (sometimes as fumarate). The rate of PEP carboxylation is relatively low (Abadie *et al.*, 2018) and so does not contribute significantly to TPU. Gauthier et al. (2010) found that amino acids made from *α*-ketoglutarate are quickly labeled by ^15^N-ammonium nitrate but not ^13^CO_2_ fed to photosynthesizing leaves indicating that the carbon for these amino acids comes from preexisting pools and so do not contribute to TPU. In Arabidopsis a significant amount of carbon is stored in the vacuole as fumarate; it is not known how much of this carbon is recent (and therefore contributes to TPU) and how much is preexisting carbon (Chia *et al.*, 2000; Pracharoenwattana *et al.*, 2010; Zell *et al.*, 2010). This is also true of sunflower (Abadie *et al.*, 2018).

## TPU and sink strength

The sink strength important in TPU limitation is different from larger scale, longer timeframe sinks such as fruit or root growth, though the two may be related. TPU is concerned with the short term, and the ability to quickly remove carbon from the Calvin-Benson cycle. Over a longer timeframe, a greater sink can be important in freeing up short-term sinks. It has been reported that defruited wheat experiences significant downregulation of photosynthesis (King *et al.*, 1967), though not all plants experience this effect (Farquhar and von Caemmerer, 1982).

In some experiments using conditions consistent with TPU, starch builds up and causes a decline in photosynthetic rate (Sasek *et al.*, 1985; Peet *et al.*, 1986; Ramonell *et al.*, 2001). However, a long-term sink which can absorb carbon, or time under reduced stress to make use of the starch, will allow the plant to recover (Sasek *et al.*, 1985; Arp, 1991).

## Adaptation of TPU

The capacity for triose phosphate utilization is not immutable. Plants grown under poor conditions are highly adaptive, and those grown under low temperature tend to have greatly elevated TPU capacity (Guy *et al.*, 1992; Sage and Kubien, 2007). This acclimation largely comes from increased expression of sucrose biosynthesis enzymes (Guy *et al.*, 1992; Hurry *et al.*, 2000). This increased capacity offsets the decreased activity of starch synthase and sucrose-phosphate synthase at low temperature and makes it less likely that the plant will be TPU limited (Cornic and Louason, 1980; Sage and Sharkey, 1987). Plants transferred to an elevated CO_2_ environment developed increased phosphate regeneration capacity, demonstrating adaptation (Sharkey *et al.*, 1988; Sage *et al.*, 1989). Plants experiencing water stress reduce their TPU capacity, possibly reflecting the reduced internal CO_2_ partial pressure that results from stomatal closure (von Caemmerer and Farquhar, 1984; Vassey and Sharkey, 1989; Cornic *et al.*, 1992). Transgenic plants overexpressing alternative oxidase cope better with water stress (Dahal *et al.*, 2014, 2015) and experience reduced negative effects on assimilation from TPU capacity. The reduced occurrence of TPU limitation in plants overexpressing the alternative oxidase was correlated with higher amounts of chloroplast ATP synthase, which might allow ATP synthesis at lower phosphate concentration. This adaptability shows that TPU will influence the metabolic investments of the plant; it will enhance the ability to handle high TP production, but only when it is required for the current output of photosynthesis.

The adaptability of TPU is important for fulfilling the role of phosphate in balancing starch synthesis and ATP synthesis (Fig. 2). Starch synthesis is highly sensitive to phosphate due to inhibition of ADP-glucose pyrophosphorylase (Preiss, 1982), and ATP synthase is kinetically sensitive to phosphate (Takizawa *et al.*, 2007). This relationship can help explain the very low partitioning of carbon into starch at low photosynthetic rate (Escobar-Gutiérrez and Gaudillère, 1997). The greatest productivity will be achieved with a fine balance. In environments less likely to be limited by TPU the plant benefits from reduced TPU capacity, which allows stromal phosphate to drop and activate starch synthesis. Conversely, in stressful environments with TPU-genic conditions, increased TPU capacity will allow better recycling of phosphate and improved ATP synthase conductivity.

**Fig 2.**
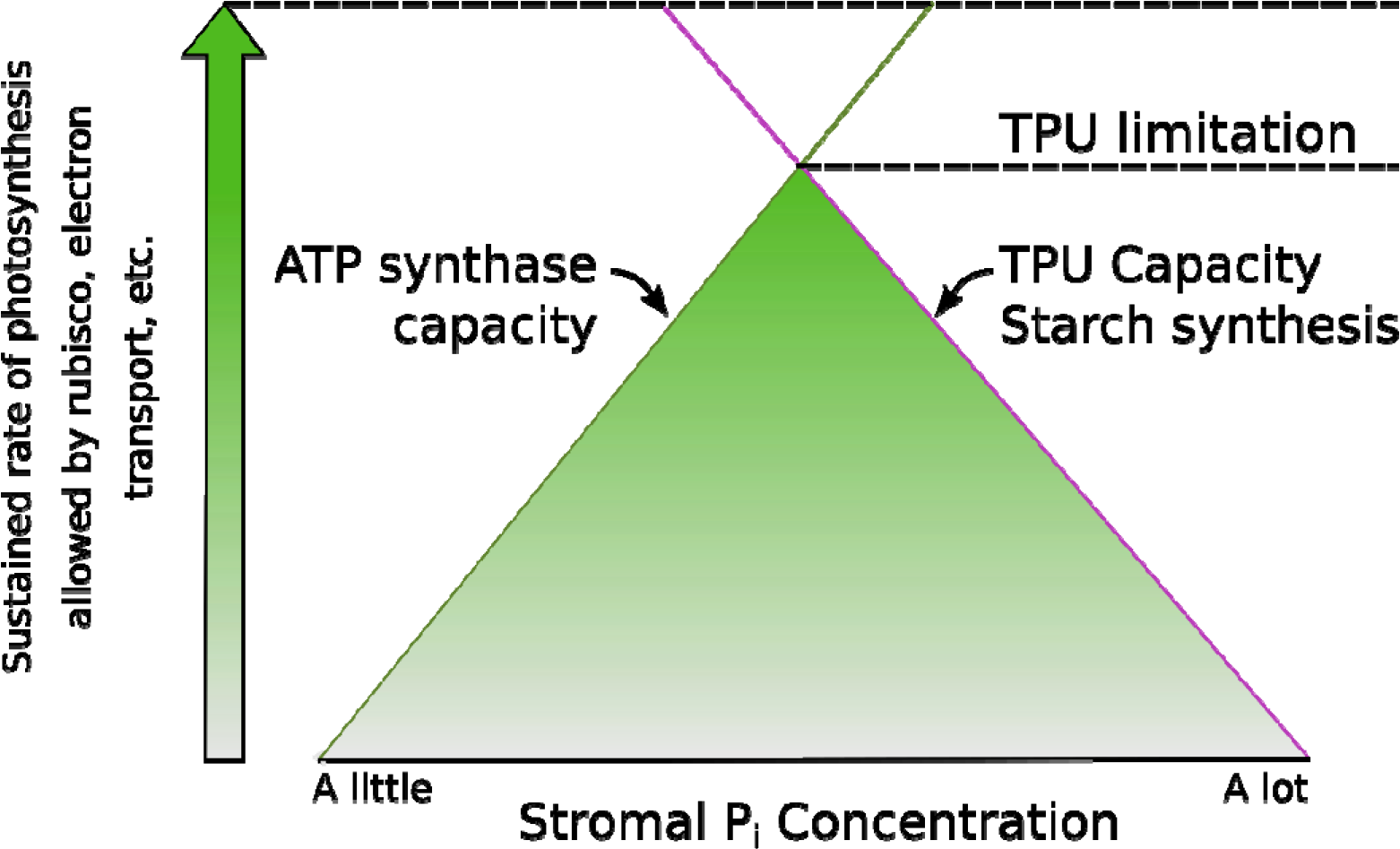
As photosynthetic rate increases, the gap between the phosphate concentration required by the ATP synthase and the phosphate concentration to inhibit starch synthesis narrows. The shapes of the responses are represented by straight lines only for simplicity. When TPU limits photosynthetic rate any increase in phosphate required for higher ATP synthase activity would inhibit starch synthesis restricting phosphate release.

## TPU and gas exchange

TPU is typically assessed from gas exchange data obtained using infrared gas analyzers to measure rates of CO_2_ uptake. Because of the usefulness of fluorescence parameters in analyzing gas exchange data, discussed later, gas exchange measurements are frequently combined with chlorophyll fluorescence. Modifying the external CO_2_ partial pressure and measuring the stomatal conductance to gas exchange by transpiration allows the calculation of the partial pressure of CO_2_ inside the leaf (*C*_*i*_) (Sharkey *et al.*, 1982). Diffusion resistance within the mesophyll will further reduce the effective partial pressure of CO_2_ resulting in the partial pressure of CO_2_ at the site of carboxylation (*C*_*c*_). This review will not include discussion of the diffusive limitations of photosynthesis. For a review of these topics, we recommend Flexas et al. (2008) and Evans et al. (2009).

The assimilation of CO_2_ (*A*) is measured by depletion of CO_2_ from air passing over the leaf. Plots of *A* against *C*_*i*_ (or better *C*_*c*_ when mesophyll conductance is known or can be estimated) can be interpreted using rubisco kinetics to predict what biochemical process is limiting assimilation. At low *C*_*i*_, assimilation is typically limited by binding kinetics of CO_2_ to rubisco, known as the rubisco limitation or C limitation. At intermediate photosynthetic rates, assimilation is typically limited by the rate of regeneration of ribulose 1,5-bisphosphate (RuBP). RuBP regeneration requires the products of the light reactions and so is frequently called J limitation. TPU limitation, sometimes called P limitation, only happens when the plant is photosynthesizing very quickly. This requirement for high photosynthetic rate may be why TPU is so hard to detect in Arabidopsis (Yang *et al.*, 2016). It is important to know the physiological differences between the three for use in fitting models because the three limitations are sometimes hard to distinguish from one another. This is especially important as many fitting models will require the user to manually select which data points are limited by each biochemical process.

By current knowledge, C_4_ plants do not suffer TPU limitation. C_4_ plants at high photosynthetic rates are interpreted to be limited by rubisco activity, and at lower rates by PEP carboxylase activity (Collatz *et al.*, 1992). The carbon pump of C_4_ metabolism makes it impossible to see the gas exchange behaviors that characterize TPU limitation.

## Physiology of TPU limitation

### Insensitivity to increasing CO_2_ and O_2_ partial pressures

TPU is characterized by a lack of, or reverse, sensitivity to oxygen partial pressure changes and CO_2_ partial pressure increases (Sharkey, 1985*a*). When photosynthesis is limited either by rubisco or RuBP regeneration, increasing CO_2_ or decreasing O_2_ would be expected to increase photosynthetic output. When TPU is limiting this is not seen, and sometimes the reverse is seen to a small extent (Ludwig and Canvin, 1971; Jolliffe and Tregunna, 1973; von Caemmerer and Farquhar, 1981). This insensitivity happens because the controlling factor is the ability of the leaf to make end products and rubisco does not need to be optimally productive to achieve maximum carbon assimilation. Increasing photorespiration means that the plant will have to perform more photosynthesis to compensate for the carbon loss, but as long as the rate of RuBP regeneration is sufficient to support the increase in both photorespiration and photosynthesis there will be no change in assimilation. (If the rate of RuBP regeneration is not sufficient to support this change, the plant would then be considered J limited). In the same manner, increasing CO_2_ further does reduce φ but the maximum carbon efflux from the Calvin-Benson cycle cannot increase and so net assimilation stays constant.

The insensitivity to oxygen is the most definitive diagnostic test to distinguish between TPU and C or J limitations. Under J limitation, assimilation is modeled as

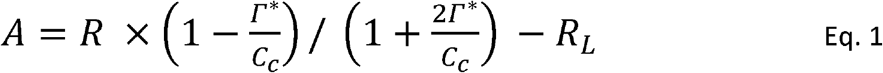

where R_*L*_ is respiration in the light (also known as *R*_*d*_), *R* is the rate of reactions of rubisco or of RuBP usage, and *Γ*\* is the rubisco CO_2_ compensation point without the effect of *R*_*L*_ (Farquhar and von Caemmerer, 1982). *Γ*\* is linearly sensitive to oxygen, and a tenfold reduction in oxygen partial pressure results in a tenfold reduction in Γ* (Forrester *et al.*, 1966; Laing, 1974). The oxygen sensitivity of assimilation while J is limiting is given in equation 2:

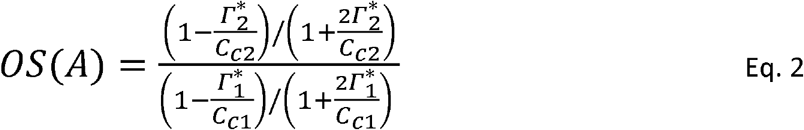

*R* from equation 1 is assumed to remain constant as it is the limiting factor in J limitation, and so it does not appear in equation 2. Equation 2 models the stimulation in photosynthetic carbon assimilation when oxygen is reduced resulting in reduced photorespiration. TPU limitation was defined by lack or reduction of this stimulation (Badger *et al.*, 1984; Sharkey, 1985a). This difference stems from the nature of the limitation. J-limited photosynthesis increases with increasing CO_2_ and decreasing O_2_ because it increases the proportion of generated RuBP used for carboxylation and reduces the CO_2_ released by photorespiration. The photosynthetic electron transport rate (*ETR*) is constant by definition. Under C limitation, changing CO_2_ or oxygen partial pressure modifies assimilation according to rubisco binding kinetics. Increased CO_2_ means more substrate, and reduced O_2_ relieves competitive inhibition resulting in higher assimilation. In C-limited photosynthesis *ETR* increases with increases in both CO_2_ and oxygen since both carboxylation and oxygenation consume RuBP. These effects are not seen at TPU limitation, as net carbon fixation is at a maximum and increases in CO_2_ or decreases in O_2_ do not increase the capacity of the plant to produce end products. Leegood and Furbank (1986) demonstrated an example of oxygen-insensitive photosynthesis in leaf discs induced by a combination of low temperature and high CO_2_ partial pressures. Feeding of phosphate restored normal oxygen sensitivity and also increased CO_2_ assimilation rate, showing that phosphate metabolism was responsible for both oxygen sensitivity and the limitation of assimilation.

### Reverse sensitivity to increasing CO_2_ and O_2_ partial pressures

While the TPU limitation offered understanding of insensitivity to increasing O_2_ and CO_2_ partial pressures, it did not immediately explain reverse sensitivity. It has long been known that oxygen inhibits photorespiration due to the competitive binding and photorespiratory CO_2_ release (Warburg, 1919; Ludwig and Canvin, 1971; McVetty and Canvin, 1981). It was therefore unexpected to find that reducing oxygen or increasing CO_2_ partial pressures could sometimes reduce the rate of CO_2_ assimilation. As photorespiration releases CO_2_, it would not make sense that altering the gas composition to favor oxygenation would result in increased carbon assimilation. Yet data dating back decades shows that once at high CO_2_, increasing CO_2_ can cause a decrease in net assimilation (Jolliffe and Tregunna, 1973; Canvin, 1978; von Caemmerer and Farquhar, 1981), and increasing O_2_ can cause an increase in net assimilation (Viil *et al.*, 1977).

Photorespiration was one key to understanding the reverse oxygen sensitivity under TPU conditions. Phosphate is temporarily released from phosphoglycolate by phosphoglycolate phosphatase before export through PLGG1 to the peroxisome. Photorespiratory metabolism of two glycolate molecules leads to re-importation of glycerate, which is phosphorylated into phosphoglyceric acid. The extra phosphate released can be used to make ATP that phosphorylates ribulose 5-phosphate to produce RuBP that will be used to accept a CO_2_, balancing the photorespiratory loss of one carbon. However, there are two amino acid intermediates in the photorespiratory pathway which could be used in the cytosol and not re-imported into the Calvin-Benson cycle. This carbon is effectively lost from RuBP, as all carbon from glycolate comes from RuBP and none of it is directly from CO_2_ fixed from the atmosphere. Photorespiratory carbon that never returns to the chloroplast was parameterized as *α*, the fraction of carbon that enters as glycolate carbon and which leaves as amino acids (Harley and Sharkey, 1991). The *α* parameter was later refined to *α*_G_ and *α*_S_, the fraction of glycolate carbon that leaves as glycine and serine respectively (Busch *et al.*, 2018). When glycine is exported instead of serine, no CO_2_ is released which would otherwise be available for refixation or diffusion out of the leaf. As these amino acids come from phosphorylated plastidic metabolites, and permanently leave the Calvin-Benson cycle, they contribute to TPU capacity. Decreasing *φ* reduces the export of glycine and serine and therefore reduces TPU capacity, explaining reverse sensitivity to CO_2_ and O_2_.

Starch synthesis is also affected by oxygen partial pressure and can contribute to severe reverse sensitivity. Beans photosynthesizing quickly then transferred to low oxygen were found to have reduced rates of starch synthesis and minimal effects on sucrose synthesis. A concurrent reduction in the ratio of glucose-6-phosphate to fructose-6-phosphate indicates inhibition of phosphoglucose isomerase (Dietz, 1985; Vassey and Sharkey, 1989). The precise mechanism of this inhibition is unclear.

### Effects on the light reactions

Elevating CO_2_ partial pressure when photosynthesis is limited by TPU will cause a decrease in *Φ*_*PS2*_. Rubisco binds CO_2_ and O_2_ competitively, meaning that an increase in CO_2_ partial pressure reduces the rate of use of RuBP for oxygenation. When TPU is controlling this does not lead to an increase in assimilation. Both carboxylation and oxygenation require ATP and NADPH, which come from electron transport. Increased CO_2_ partial pressure therefore causes reduced velocity of rubisco and an overall reduction in electron transport requirements (Stitt, 1986; Sharkey *et al.*, 1988; Stitt and Grosse, 1988). Regulatory processes lead to reduced *Φ*_*PS2*_, a phenomenon which can be useful in discriminating TPU limitation from combined gas exchange and fluorescence data (Fig. 3).

**Fig 3.**
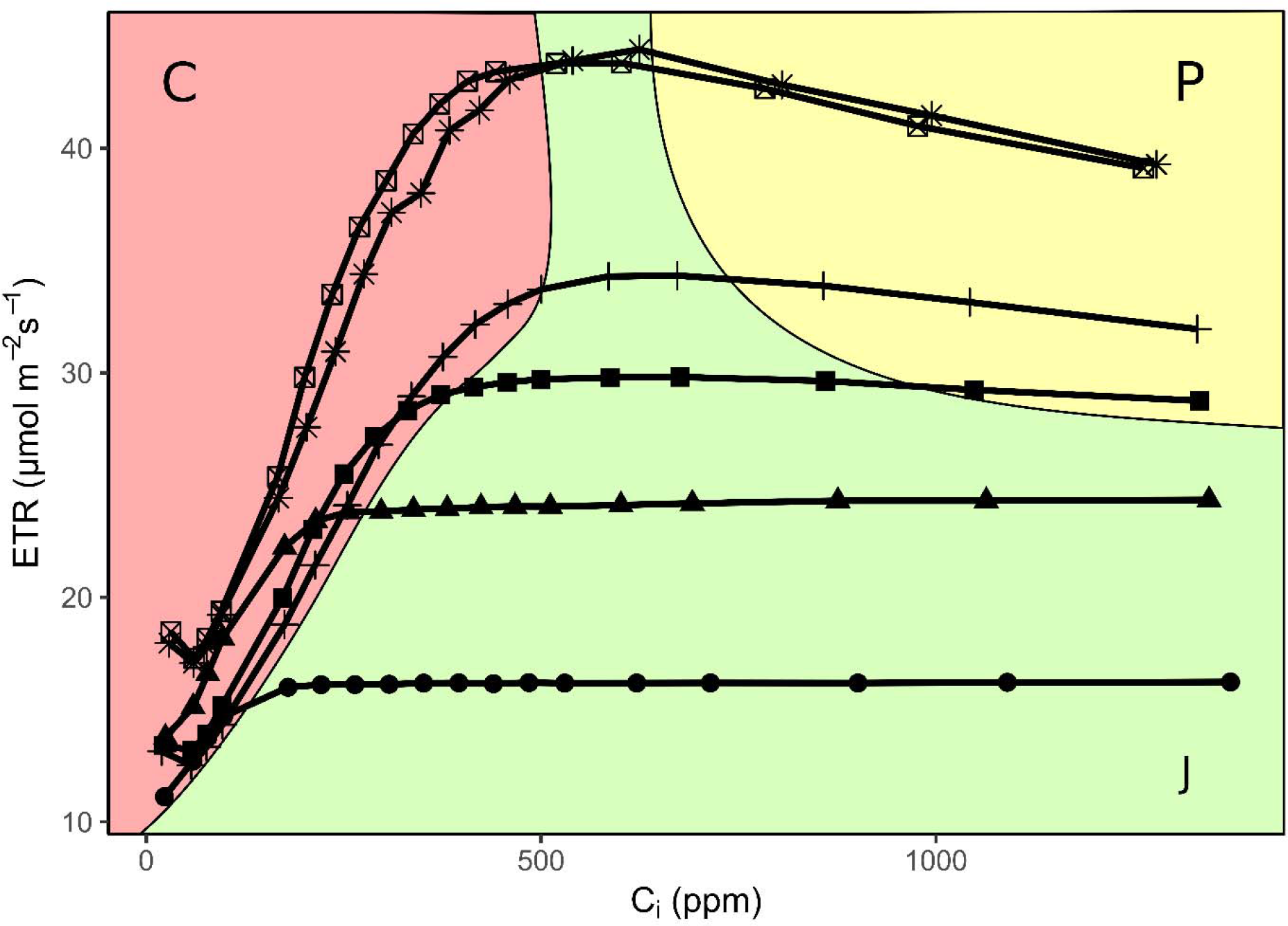
The decline in electron transport rate is diagnostic of TPU limitation. From combined gas exchange and fluorescence data in *A/C*_*i*_ curves of *Nicotiana benthamiana* at varying light intensity and 35°C. At low CO_2_, plants are limited by rubisco activity (C limitation, red), characterized by a sharp upwards slope of both A and *ETR* with increasing CO_2_. When light is insufficient, plants will be limited by the rate of RuBP regeneration (J limitation, green), characterized by a flat slope of *ETR* with increasing CO_2_. Only when the plant has ample CO_2_ and electron transport will TPU limitation (P, yellow) be seen, characterized by a decline in *ETR* with increasing CO_2_. *ETR* is calculated from fluorescence-derived *Φ*_*PS2*_. Light intensity (µmol m^−1^ s^−1^): •- 250, ▴- 400, ▀- 550, +- 750, ⊠- 1000, *- 1500.

There are effects on the kinetics of the light reactions that happen concurrently with reduction of electron transport rate. Proton conductivity across the thylakoid membrane goes down under TPU limitation (Takizawa *et al.*, 2008; Kiirats *et al.*, 2009; Yang *et al.*, 2016). It is proposed that this kinetic change occurs because of a reduced pool of available phosphate in the stroma which reduces the rate of ATP synthase. The *K*_*m*_ of chloroplast ATP synthase for phosphate has been measured at 0.2-1 mM (Selman-Reimer *et al.*, 1981; Grotjohann and Gräber, 2002). Stromal phosphate concentration during feedback conditions is estimated to be between 0-1.7 mM depending on how much phosphate is assumed to be free (Sharkey and Vanderveer, 1989), so it is reasonable to suggest that the phosphate concentration may drop below the *k*_*m*_ of ATP synthase. Joint with a decrease in ATP synthase conductivity is an increase in proton-motive force (*PMF*). The energy needed to make ATP will depend on the concentration of phosphate (Eq. 3).

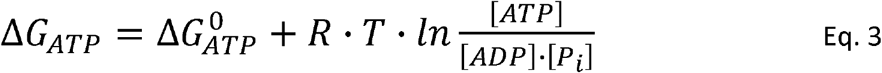

As the effective [P_i_] declines, *ΔG*_*ATP*_ will increase, requiring a greater *PMF* for an equivalent rate of ATP synthesis. Increased *PMF* leads to controls on electron transport through energy-dependent exciton quenching (*q*_*E*_) reducing energy arrival at P680 or reduction in the rate of electron flow at the cytochrome *b*_*6*_*f* complex, leading to reduced rates of electron transport (Kramer and Crofts, 1996; Owens, 1996). While phosphate seems to play a role in linking the light reactions and the Calvin-Benson cycle, it is less clear what other molecular mechanisms may be important. It is likely that we do not yet know some important regulatory components that control *ETR* when TPU limits the rate of photosynthesis.

### Temperature sensitivity

Photosynthesis under TPU limitation is highly temperature sensitive. Rubisco-limited photosynthesis is largely unaffected by temperature because of increasing rates of both respiration and photorespiration and their effect on the rubisco CO_2_ compensation point (Sage and Kubien, 2007). J-limited photosynthesis may be affected by temperature due to the temperature dependence of the maximum rate of electron transport (*J*_*max*_) (June *et al.*, 2004; Cen and Sage, 2005; Sage and Kubien, 2007). TPU conditions are the most affected by temperature (Yang *et al.*, 2016) perhaps because of the strong temperature sensitivity of sucrose-phosphate synthase (Stitt and Grosse, 1988; Leegood and Edwards, 1996). Other enzymes implicated in TPU limitation are also temperature sensitive, such as nitrate reductase (Leegood and Edwards, 1996; Busch *et al.*, 2018). Because of the different ways by which temperature affects the three limitations, the conditions in which they appear changes with temperature. Notably, at temperature lower than growth conditions the plant is significantly more likely to become TPU limited (Stitt, 1986; Sage and Sharkey, 1987; Labate and Leegood, 1988). Labate and Leegood (1988) demonstrated a temperature-sensitive increase in photosynthesis from phosphate feeding. Leaf discs floated on a solution containing phosphate at 25°C saw a marginal reduction in assimilation. However, discs fed phosphate at 10°C experienced significant photosynthetic gains, indicating that reduced temperatures result in greater limitation of photosynthesis by TPU (Fig. 4).

**Fig 4.**
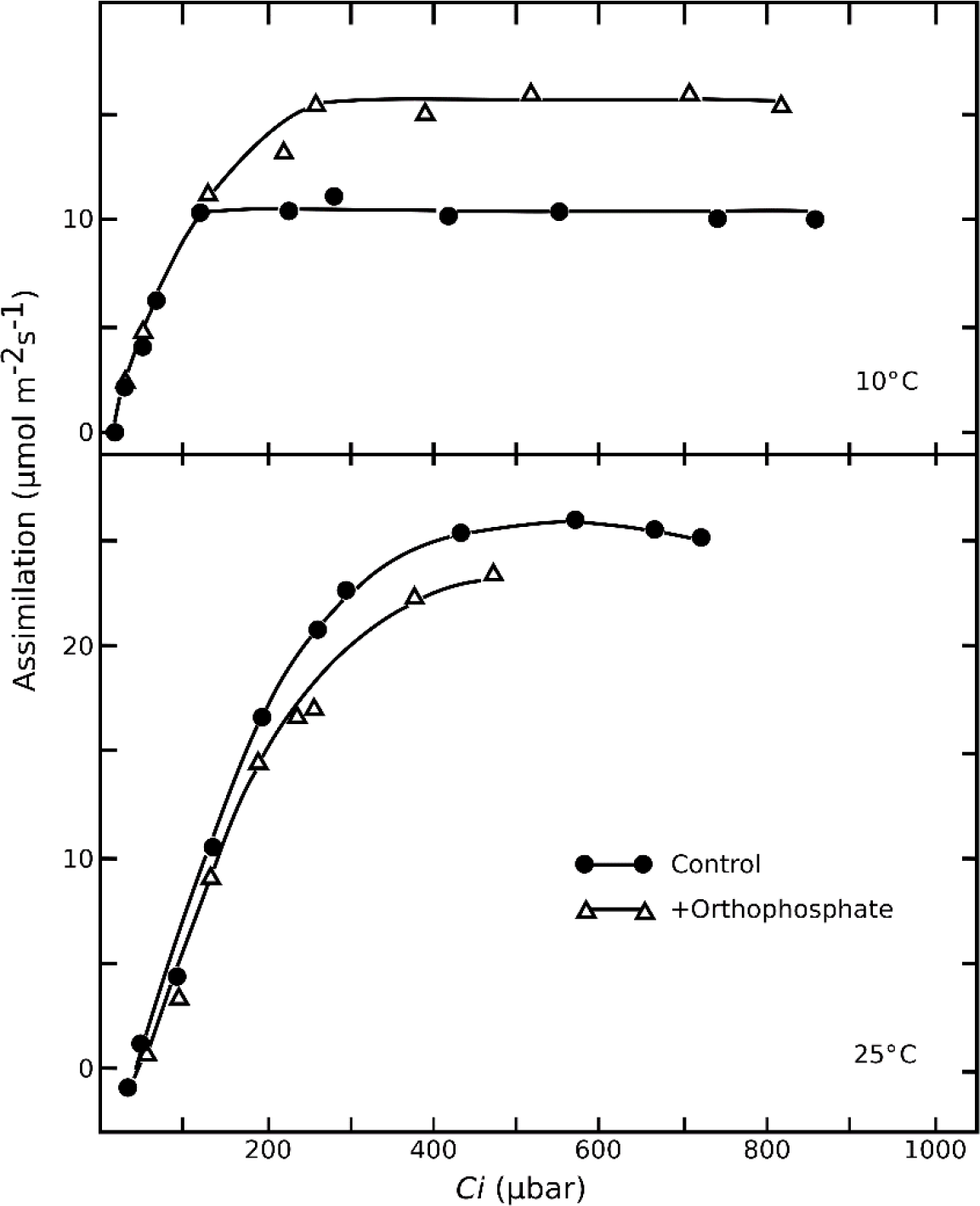
Rate of CO_2_ assimilation of barley versus *C*_*i*_ in 10°C (top) and 25°C (bottom) with and without the addition of phosphate. A temperature-dependent increase in photosynthetic assimilation is observed upon addition of phosphate. Re-drawn from Labate & Leegood 1988.

### TPU and plant nutrition

TPU is often incorrectly interpreted as a nutritional deficiency. It is true that plants transferred to media without any phosphate experience significant reduction in photosynthetic capacity (Brooks, 1986; Foyer and Spencer, 1986). However, less dramatic differences in phosphate nutrition result in relatively small changes in photosynthetic rate. This is due to the vacuole buffering phosphate concentration in the rest of the cell on a hours timescale (Rebeille *et al.*, 1983; Woodrow *et al.*, 1984). Under increased or decreased phosphate nutrition, large changes in vacuolar phosphate concentration are seen, but only relatively small changes are seen in plastidic phosphate concentration (Rebeille *et al.*, 1983; Foyer and Spencer, 1986). Plants grown with different phosphate nutrition are therefore not significantly more or less likely to experience TPU limitation. Ellsworth et al. (2015) showed a survey of Australian plants growing in the wild with varying phosphate availability were adapted to their environment, and TPU was more likely at ***high*** phosphate nutrition. Furthermore, TPU can only be seen when the plant is photosynthesizing very quickly, which usually cannot be seen if the plant is nutritionally deprived. Plants with reduced nitrogen were not capable of photosynthesizing quickly enough to reach TPU limitation (Sage *et al.*, 1990).

### Oscillations

Oscillations are a common side-effect of TPU limitation (Ogawa, 1982; Sivak and Walker, 1986, 1987). They are typically seen after a perturbation in the conditions of a plant in high photosynthetic conditions. Oscillations include tandem changes in carbon assimilation and fluorescent parameters, indicating simultaneous changes in both the light reactions and the Calvin-Benson cycle (Ogawa, 1982; Walker *et al.*, 1983; Peterson *et al.*, 1988; Stitt and Grosse, 1988). The amplitude of oscillations can increase with conditions that further exacerbate TPU limitation, such as low temperature or low O_2_ (Peterson *et al.*, 1988; Stitt and Grosse, 1988). Oscillations showed a significant impact on organic phosphates and their relevant ratios, notably large initial spikes in PGA, reduction in RuBP and ATP pools (Sharkey *et al.*, 1986b; Sage *et al.*, 1988; Stitt and Grosse, 1988; Laisk *et al.*, 1991).

A number of models have been produced to explain oscillations. The most significant theory is that there is a delay in activation of sucrose synthesis after a photosynthetic increase which causes oscillations (Laisk and Walker, 1986). This delay may be due to fructose-2,6-bisphosphate levels, a compound with numerous regulatory roles including inhibition of cytosolic fructose bisphosphatase activity (Stitt *et al.*, 1984; Laisk and Eichelmann, 1989; Laisk *et al.*, 1989). An additional interpretation of these oscillations has been proposed originating from the light reactions, with damping caused by a slow leak of protons across the thylakoid membrane (Kocks and Ross, 1995).

## Modeling

TPU models have seen some changes recently. Original models that account for triose phosphate usage relied on simple stoichiometry (Sharkey, 1985*b*):

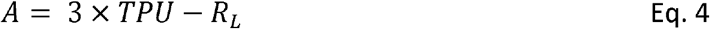

Under this model photosynthetic assimilation would exactly equal the rate of carbon export from the Calvin-Benson cycle for starch and sucrose synthesis. However, this model did not account for reverse sensitivity of assimilation to oxygen or CO_2_ frequently observed, and the model’s predictive power declined at increasing oxygen partial pressures. The model also describes all carbon export as triose phosphate usage, which is not directly true. Any carbon that leaves the Calvin-Benson cycle and is dephosphorylated will contribute to the maximum assimilation rate. While all carbon in the Calvin-Benson cycle derives from TP, some of the end products are made from Calvin-Benson cycle intermediates other than TPs. Despite this, the simple model has some advantages. It requires no estimation of *R*_*L*_, mesophyll conductance (*g*_*m*_) or *Γ*\*. These parameters are used to fit the other limitations and for other models of TPU. These three parameters are currently impossible to directly measure, and there is some debate about our ability to accurately fit them and the constancy of these parameters.

The more recent and complex model for TPU incorporates parameters for photorespiratory amino acid release. From Busch et al. (2018):

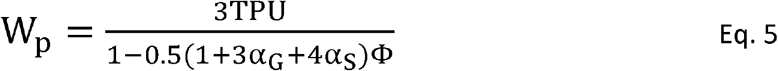

where *W*_*p*_ is the rate of carboxylation when limited by phosphate metabolism. The denominator in equation 5 adjusts the rate of TPU by the fraction of carbon that is exported as triose phosphates, rather than exported during photorespiration due to: CO_2_ loss during recombination of glycine to serine; direct export of glycine; or direct export of serine (respectively, terms in parentheses). As one carbon out of four is lost as CO_2_ in the formation of serine, *α*_S_ cannot be greater than 0.75. These parameters have an impact on curve fitting for all limitations, and it notably explains the common, minor reverse sensitivity of TPU-limited assimilation to oxygen. For comparison of equations 4 and 5, *W* must be adjusted for respiratory carbon loss:

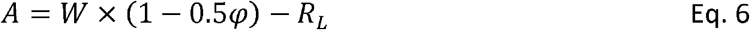

In light of equation 6, if *α*_G_ and *α*_S_ are zero, equations 4 and 5 are identical. Unlike the simple model, equation 5 requires knowledge of the relative rate of photorespiration, and therefore relies on fitting for *Γ*\*. There is also little signal to differentiate *α*_S_ and *α*_G_, which can make fitting these two parameters separately by gas exchange somewhat challenging.

Some other physiological features can be seen in the models. Decreasing the partial pressure of oxygen will decrease *φ*, and in addition to decreasing *A* this will also decrease *W* per equation 5. An oxygen decrease therefore results in reduction of both the velocity of oxygenation and the velocity of carboxylation, a net decrease in *R*, and the concomitant decrease in *Φ*_*PS2*_ as described above.

Current modeling software is available, with varying amounts of parameters to fit. Sharkey (2016) presented an excel tool which allows picking of points from *A/Ci* curves, with options to fit *R*_*L*_, *g*_*m*_, and *α*_*G*_ and *α*_*S*_. Bellasio, et al. (2016) provide an Excel tool that uses combined gas exchange and fluorescence to fit *R*_*L*_, *g*_*m*_, *J*_*max*_, *V*_*cmax*_, *Γ*\* and rubisco specificity for CO_2_ versus oxygen (*S*_*c/o*_), but not *α*; much of the basis of this fitting are also discussed by Yin et al. (2009). Dubois et al. (2007) provide a SAS program which allows fitting of *R*_*L*_, *g*_*m*_, *J*_*max*_, *V*_*cmax*_, *Γ*\* and *S*_*c/o*_, and *α*. Moualeu-Ngangue et al. (2017) propose to improve the Dubois fitting by reducing the number of assumptions made, though they do not fit *α*. It should be noted that no current model attempts to incorporate other carbon sinks.

## Environmental impact

The changing climate resulting in large measure from increasing CO_2_, has the potential to affect the frequency and severity of TPU limitations to photosynthesis. Since this syndrome occurs when carbon fixation and light capture are going faster than end products can be made, increasing CO_2_ should increase the occurrence of TPU. However, because TPU is stimulated by increasing temperature, there could be a reduction in the occurrence of TPU limitation in the future. It is hard to predict which effect will dominate, and whether TPU limitation will be observed more or less frequently based on climate change predictions. However, beyond the short-term effects of temperature and CO_2_ it is important to consider how the plant responds when it is TPU-limited. Generally, plants growing in elevated CO_2_ show less propensity for TPU limitation but because they have reduced capacity for other processes in photosynthesis (Sage *et al.*, 1989). We hypothesize that understanding TPU will help in predicting acclimation responses of plants to increasing atmospheric CO_2_.

It is often found that TPU-limitation occurs whenever photosynthesis is stimulated to be about 20% higher than was occurring in the plant under natural conditions (Yang *et al.*, 2016). Increasing CO_2_, decreasing oxygen, or lowering the temperature usually allows TPU-limitation to be observed. In a large study of published A/*C*_*i*_ curves Wullschleger (1993) found 23 cases (out of 109) where investigators reported TPU limitations. It is likely that the phenomenon is observed but not recognized much more often. For example, a curve presented in Wullschleger et al. (Figure 1B, taken from Ireland *et al.*, 1988) shows evidence of TPU-limitation but this was not one of the 26 instances of TPU limitation cited. It is common for the TPU limitation to be ignored even when it is clearly evident in data.

Since the components of photosynthesis must all work in concert and in strict stoichiometry, it is not surprising that there might be a relationship between *V*_*cmax*_ and TPU capacity. This has been invoked in global models of photosynthesis although many models do not include TPU. Lombardozzi et al. (2018) used several estimates of the ratio of *V*_*cmax*_ and TPU capacity and concluded that current global models may overestimate how much CO_2_ will be fixed by plants in the future because TPU-limitations, or adjustments to avoid TPU limitation will reduce photosynthetic capacity. It is important to realize that even though plants growing in elevated CO_2_ do not show TPU-limitation, TPU still may be setting an upper bound and that plants adjust other capacities to keep below the upper bound of TPU because TPU can cause damage.

## Conclusions

TPU is a metabolic condition that incorporates numerous signals to reflect the state of photosynthesis across the whole cell. Most metabolites in the chloroplast are phosphorylated, and as such phosphate is a good representation of the metabolic state of the chloroplast. Phosphate is linked through the cytosol, where sucrose synthesis takes place, and thus phosphate represents photosynthetic state across all chloroplasts. Phosphate concentrations and the ability to regenerate them are carefully regulated so that TPU capacity is just slightly above the highest rate of photosynthesis the plant is likely to achieve. A low phosphate level naturally signals to the other processes that photosynthesis is running very fast, kinetically controls the ATP synthase, and leads to downstream effects on photosynthesis by accumulation of *PMF* and the engaging of *q*_E_. The reduction in phosphate signals the plant to build up starch by relieving phosphate inhibition of ADP-glucose pyrophosphorylase (Preiss, 1982). Adjustment in TPU capacity allows plants which are photosynthesizing slowly to lower their phosphate regeneration and helps produce starch; conversely, it allows plants which are photosynthesizing quickly to raise their phosphate regeneration and helps produce ATP. In this way, TPU sets the spans on expected photosynthesis. We believe that the gas exchange behavior in TPU conditions reflects a number of important regulatory features. Yet, the role of TPU as regulation is relatively unexplored.

A number of misconceptions cloud the field in regards to TPU. Even the term “TPU” can now be seen not to be wholly accurate. It largely describes phosphate metabolism, but not all effects on carbon metabolism related to phosphate can be accurately described as triose phosphate usage. At steady state, there are other sources of phosphate release that contribute to the assimilation cap. Amino acid release from photorespiration, MEP and shikimate pathways, and other short-term carbon sinks will all contribute to the maximal assimilation rate when TPU limited. An alternative view is that all Calvin-Benson cycle exports are downstream of TP, and thus constitute a form of TPU. The specific terminology and nuance are less important than the total understanding, which is that TPU limitation is the result of insufficient speed of carbon export into short-term sinks. Other, longer term sinks, while they will be important for the overall physiology of the plant, will not be discernible in gas exchange measurements.

TPU is also a somewhat misleading under steady state, because the plant will not remain at feedback limitation. Maintaining feedback limitation is unhealthy for the plant due to risk of oxidative stress from photosystem oxidation (Pammenter *et al.*, 1993). Electron transport quenching and deactivation of rubisco lead to an overall slowing of photosynthesis lower than feedback, eventually reaching a steady state with assimilation rate based on the rate of carbon export. Excess assimilation when already low on phosphate would further deprive ATP synthase of phosphate it needs. Contrary to what one might expect given the term “TPU limitation,” triose phosphates do not necessarily need to build up during TPU, though phosphate levels should be low (Sharkey and Vanderveer, 1989). This is why plants can be drained of phosphate via mannose or deoxyglucose feeding and be TPU limited (Herold and Lewis, 1977; Herold, 1980; Sivak and Walker, 1986). It is the relationship between the need for phosphate for ATP synthase and the phosphate sensitivity of starch synthase that results in TPU (Herold, 1980).

## Acknowledgements

This research was funded by U.S. Department of Energy Grant DE-FG02-91ER2002 (T.D.S. and A.M.M.). A.M.M. is partially supported by a fellowship from Michigan State University under the Training Program in Plant Biotechnology for Health and Sustainability (T32-GM110523). Partial salary support for T.D.S. came from Michigan AgBioResearch.

Figure 4 Adapted by permission from Springer: Springer Planta. Limitation of photosynthesis by changes in temperature, Labate, CA, Leegood, RC, Copyright 1988.

## Abbreviations

*A*: Net assimilation of carbon, or, when negative, net respiration of carbon
*C*_*c*_: Partial pressure of CO_2_ at the site of carboxylation
*C_i_*: Partial pressure of CO_2_ inside the leaf
DHAP: Dihydroxyacetone phosphate
E4P: Erythrose-4-phosphate
*ETR*: Photosynthetic electron transport rate
FBPase: Fructose bisphosphatase
GAP: Glyceraldehyde phosphate
*g*_*m*_: Mesophyll conductance to CO_2_
*J*_*max*_: Maximum rate of electron transport under infinite light
MEP: Methylerithrytol phosphate
PEP: Phosphoenolpyruvate
PGA: Phosphoglyceric acid
PMF: Proton-motive force
PPT: Phosphoenolpyruvate/phosphate translocator
*q*_*E*_: Energy dependent quenching of electron transport
*R*_*L*_: Rate of RuBP consumption
RL: Rate of respiration in the light
RuBP: Ribulose 1,5-bisphosphate
*S*_*c/o*_: Specificity of rubisco for CO_2_ versus oxygen
TP: Triose phosphate
TPT: Triose phosphate/phosphate antiporter
TPU: Triose phosphate utilization
*V*_*cmax*_: Maximum velocity of carboxylation
W: Rate of carboxylation
*α*: The fraction of glycolate carbon which leaves the CB cycle as amino acids
*α_G_*: The fraction of glycolate carbon which leaves the CB cycle as glycine
*α_S_*: The fraction of glycolate carbon which leaves the CB cycle as serine
*Γ*\*: The rubisco CO_2_ compensation point ignoring the effect of *R*_*L*_
*φ*: The ratio of oxygenations to carboxylations
*Φ*_*PS2*_: Quantum yield of photosystem 2.

